# Mitochondrial genomes of *Columbicola* feather lice are highly fragmented, indicating repeated evolution of minicircle-type genomes in parasitic lice

**DOI:** 10.1101/668178

**Authors:** Andrew D. Sweet, Kevin P. Johnson, Stephen L. Cameron

## Abstract

Most animals have a conserved mitochondrial genome structure composed of a single chromosome. However, some organisms have their mitochondrial genes separated on several smaller circular or linear chromosomes. Highly fragmented circular chromosomes (“minicircles”) are especially prevalent in parasitic lice (Insecta: Phthiraptera), with 16 species known to have between 9 and 20 mitochondrial minicircles per genome. All of these species belong to the same clade (mammalian lice), suggesting a single origin of drastic fragmentation. Nevertheless, other work indicates a lesser degree of fragmentation (2-3 chromosomes/genome) is present in some avian feather lice (Ischnocera: Philopteridae). In this study, we tested for minicircles in four species of the feather louse genus *Columbicola* (Philopteridae). Using whole genome shotgun sequence data, we applied three different bioinformatic approaches for assembling the *Columbicola* mitochondrial genome. We further confirmed these approaches by assembling the mitochondrial genome of *Pediculus humanus* from shotgun sequencing reads, a species known to have minicircles. All three methods indicated *Columbicola* spp. genomes are highly fragmented into 15-17 minicircles between 1,119 and 3,173 bp in length, with 1-4 genes per minicircle. Subsequent annotation of the minicircles indicated that tRNA arrangements of minicircles varied substantially between species. These mitochondrial minicircles for species of *Columbicola* represent the first feather lice (Philopteridae) for which minicircles have been found in a full mitochondrial genome assembly. Combined with recent phylogenetic studies of parasitic lice, our results provide strong evidence that highly fragmented mitochondrial genomes, which are otherwise rare across the Tree of Life, evolved multiple times within parasitic lice.

## INTRODUCTION

Mitochondria are crucial for the survival of most eukaryotic cells and organisms (although see Karnkowska et al. 2016). Because of this important function, the mitochondrial genome has a very stable structure across deep evolutionary scales. The overwhelming majority of metazoans have 37 mitochondrial genes on a single, ∼16,000 bp circular chromosome: 13 protein-coding genes (PCGs), two ribosomal RNAs (rRNAs), and 22 transfer RNAs (tRNAs) (Wallace 1982, Wolstenholme 1992, Boore 1999, Chen and Butow 2005). Significant deviations from this established genomic architecture can have dire consequences for the affected organism. For example, deletion within mitochondrial DNA in humans has been associated with several diseases, including Parkinson’s and Kearns-Sayre syndrome (Zeviani et al. 1988, Bender et al. 2006). Likewise, mitochondrial genome fragmentation has been linked to cell death, disease (e.g., cardiomyopathy), and aging (Yoneda et al. 1995, Hayakawa et al. 1996, Kajander et al. 2000, Bender et al. 2006).

Although abnormal mitochondrial genome architecture in animals is often a pathological state, several different taxa are known to possess multiple mitochondrial chromosomes under normal conditions. These fragmented mitochondria have seemingly arisen multiple times across the Tree of Life (Smith and Keeling 2015). Parasitic nematodes in the genus *Globodera* have at least 6 circular chromosomes, each of which differs in gene content and order (Gibson et al. 2007a, b). There is also evidence of recombination between the different chromosomes in *Globodera*. Fragmented mitochondrial genomes have also been identified in several insect genera including the booklouse *Liposcelis* with 2-3 chromosomes (Wei et al. 2012, Shi et al. 2016) and the thrips *Scirtothrips* with 2 chromosomes (Dickey et al. 2015). Mitochondrial fragments are not always circular, although cases of linear mitochondrial chromosomes are only known from outside of Bilateria (Bendich 1993). For example, the unicellular ichyosporean *Amoebidium parasiticum* has a large (>200 kbp) mitochondrial genome consisting of several hundred linear chromosomes (Burger et al. 2003), the box jellyfish *Alatina moseri* has 8 linear chromosomes (Smith et al. 2011), and the Hydrozoa *Hydra magnipapillata* has two linear chromosomes (Voigt et al. 2008).

The greatest known abundance of fragmented mitochondrial genomes within a single group is in the parasitic lice (Insecta: Phthiraptera), a group comprising around 5,000 species. Many species of lice have a single mitochondrial chromosome, although in all species studied to date, the gene order is rearranged relative to the inferred ancestral insect mitochondrial genome (Shao et al. 2001, Price et al. 2003, Covacin et al. 2006, Cameron et al. 2007, 2011). However, 16 species of lice have, to date, been shown to have highly fragmented mitochondrial genomes with their genes divided across 9 to 20 “minicircles” depending on the species (Jiang et al. 2013, Dong et al. 2014a, b, Herd et al. 2015, Shao et al. 2015, 2017). The first louse species shown to have minicircles was the human louse, *Pediculus humanus* (Shao et al. 2009, 2012). There are at least 20 mitochondrial chromosomes each between 3-4 kbp long in *P. humanus*. Each circle has a distinct coding region containing an average of 2 genes, but there is also a non-coding control region (CR) present in every circle, composed largely of motifs conserved between different minicircles. This region includes the D-loop which is involved in DNA replication and the initiation of transcription (Clayton 1982, Aquadro and Greenberg 1983). Other louse taxa with minicircles vary in their number of chromosomes (between 9 and 20), but the circles have a similar structure to those found in *P. humanus*: a few thousand base pairs in length, with variable coding regions and conserved CRs. The mechanism of mitochondrial genome fragmentation in lice is unknown, although Cameron et al. (2011) reported the gene for the nuclear-encoded mitochondrial single-stranded DNA-binding protein (mtSSB) is either missing or non-functional in *Pediculus*. Because mtSSB is involved in mitochondrial replication, the lack of this protein could inhibit replication of a normal, single 16kbp+ long chromosome, whereas replication of multiple, smaller chromosomes could proceed without mtSSB (Pestryakov and Lavrik 2008, Ruhanen et al. 2010).

Complete sets of mitochondrial minicircles in lice have only been found in mammalian sucking lice (Anoplura), elephant and warthog lice (Rhyncophthirina), and mammalian chewing lice of the family Trichodectidae. Together these lice form a highly-supported monophyletic group (Johnson et al. 2018), therefore implying mitochondrial fragmentation evolved once in parasitic lice (Song et al. 2019). Cameron et al. (2011) found evidence of mitochondrial genome fragmentation in three species of avian feather lice (Philopteridae), although these were only partial genomes, with only one chromosome sequenced and 6-19 genes recovered on a chromosome per louse species. These partial, fragmented mitochondrial genomes from avian lice differed substantially from those found in mammal lice. In particular, the avian louse genomes are typically larger, contain more protein-coding or ribosomal RNA genes (3-7 PCG/rRNA genes in avian lice, 0-2 in mammal lice), and lack a large (1000bp+) CR with conserved structural motifs. The fragmented mitochondrial genomes from avian lice reported by Cameron et al. (2011) more closely resembled those found in groups outside of lice (e.g. *Lipsoscelis*, *Scirtothrips*, and *Globodera*) than those found in mammalian lice. For this reason, Cameron et al. (2011) proposed definitions for the different mitochondrial genome fragmentation “types” found in avian versus mammalian lice. This distinction was emphasized by Song et al. (2019) in their proposed name Mitodivisia, the monophyletic clade composed of the three mammal-associated louse lineages, whose diagnostic feature was the minicircle form of extensive mitochondrial fragmentation. The extent of mitochondrial fragmentation in avian lice, and how similar their genome structures are to those found in mammalian lice, has yet to be evaluated.

Here, we test for mitochondrial minicircles in avian feather lice by focusing on *Columbicola*, a representative of one of the three main clades within Philopteridae (Johnson et al. 2018), and the only one for which complete mitochondrial genomes have yet to be sequenced. Species of *Columbicola* parasitize pigeons and doves and are specialized to live between feather barbs in wing feathers of their hosts, although they eat downy feathers near the host’s body (Nelson and Murray 1971, Marshall 1981, Clayton et al. 1999). These lice occur throughout the range of their host group, which includes every continent except Antarctica (Price et al. 2003). We have several motives for targeting this genus of feather lice. First, previous attempts to sequence and assemble the mitochondrial genome of *Columbicola* have been unsuccessful using traditional methods such as long PCR, suggesting the architecture of the genome is more complicated than a typical single chromosome (pers. obs.). Second, uncovering the mitochondrial genome structure of *Columbicola* would be highly informative for understanding mitochondrial evolution in parasitic lice. The phylogenomic analysis of Johnson et al. (2018) recovered *Columbicola* on a relatively long branch representing one of three main clades in Philopteridae. However, *Columbicola* is not the earliest diverging lineage in the family; a clade including *Bothriometopus*, a genus with a full-sized, single-chromosome mt genome (Cameron et al. 2007) is the sister group to the rest of Philopteridae (Johnson et al. 2018). The remaining Philopteridae form a clade sister to that represented by *Columbicola*, and includes multiple genera documented to possess full-sized, single-chromosome mt genomes (*Campanulotes*: Covacin et al. 2006; *Ibidoecous* & *Coloceras*: Cameron et al. 2011; *Falcolipeurus*: Song et al. 2019). Based on this phylogeny, *Columbicola* is therefore nested within clades of lice which possess single chromosome mitochondrial genomes. If *Columbicola* does not have a single chromosome, this would definitively support an independent origin of fragmentation (c.f. the fragmented genomes described by Cameron et al. 2011 which are partial and thus may not accurately represent genome structure in these species). Third, there is a well-supported and well-sampled phylogeny of the genus *Columbicola* from Boyd et al. (2017), which allows us to choose species that accurately represent the evolutionary diversity of the group. Finally, *Columbicola* is a model system in host-parasite ecology and evolution (Clayton and Johnson 2003, Johnson et al. 2007, Bush and Malenke 2008, Bush et al. 2010, Clayton et al. 2016). Over two decades of experimental work on the genus provides us with the most abundant knowledge-base for any group of feather lice. Previous studies of mitochondrial fragmentation in lice have related the phenomena to a variety of life-history traits shared by the taxa which possess them (e.g. life-cycle length or use of mammal hosts; Dong et al. 2014b; Song et al. 2019). Investigating mitochondrial fragmentation in a feather louse genus with well-known ecology allows us to build on and contextualize these previous hypotheses.

In this study, we use existing and novel genome sequence data to assemble the mitochondrial genomes of four *Columbicola* species that represent major clades in the genus. We utilize three different bioinformatic methods for assembling mitochondrial genomes, and then annotate the resulting sequences to test for fragmentation and rearrangements within the genus. Finally, we then confirm the reliability of our bioinformatic approach by assembling the fragmented mitochondrial genome of *Pediculus humanus*, for which mt genome structure has been previously been verified with PCR and Southern blot methods.

## MATERIALS AND METHODS

### Whole genome sequence data

We obtained whole genome shotgun Illumina sequencing reads from the NCBI SRA database for *Columbicola macrourae* ex. *Zenaida macroura* (SRR3161953), *C. passerinae* ex. *Columbina picui* (SRR3161931; hereafter “*C. passerinae* 1” following Boyd et al. (2017)), and *C. passerinae* ex. *Columbina cruziana* (SRR3161930; hereafter “*C. passerinae* 2”). We also generated new sequence data for a specimen of *C. columbae* from the feral rock pigeon (*Columba livia*). These species represent 3 of the 4 major clades identified within *Columbicola* by Boyd et al. (2017) and thus are a good representation of this speciose genus (90 described species; Bush et al. 2009). *Columbicola passerinae* is a known complex of cryptic species parasitizing New World ground-doves (Johnson et al. 2007). Although they are currently described as a single species based on morphology, individuals from *C. passerinae* form two well-defined clades structured according to mitochondrial (*cox1*) and genomic-level nuclear sequence data, biogeography, and host association (Sweet and Johnson 2016, 2018). This complex is therefore a useful system for comparing mitochondrial architecture between two very closely related species.

Each specimen was photographed as a voucher and total genomic DNA was extracted from individual lice using a Qiagen QIAamp DNA Micro Kit (Qiagen, Valencia, CA, USA) with a 48-hour incubation period at 55°C in proteinase K. The resulting DNA was quantified with a Qubit 3.0 Fluorometer (Invitrogen, Carlsbad, CA, USA) using the standard protocol, fragmented with sonication for a mean insert size of 400bp, and prepared for sequencing with a Hyper Library Preparation Kit (Kapa Biosystems, Wilmington, MA, USA). Paired-end sequencing was then performed with 151 cycles in a single lane on an Illumina NovaSeq 6000 using a NovaSeq S4 reagent kit.

For both SRA and newly generated sequence data, we trimmed adapters and filtered reads following Boyd et al. (2017). We removed duplicate read pairs with fastqSplitDups.py (https://github.com/McIntyre-Lab/mcscript), removed the first 5 nt from the 5’-end of de-duplicated reads using Fastx_clipper v.0.0.14 (http://hannonlab.cshl.edu/fastx_toolkit/), removed bases with a phred quality score <28 from the 3’-end using a 1 nt window in Fastx_trimmer v.0.014, and after trimming removed reads <75 nt. We confirmed adapter and quality trimming by running the trimmed libraries through FastQC v.0.11.7 (https://www.bioinformatics.babraham.ac.uk/projects/fastqc/).

### Mitochondrial genome assembly of *Columbicola*

We used three different approaches to assemble the mitochondrial genomes of *Columbicola.* These three approaches also served as lines of evidence that the *Columbicola* mt genome is split up into several circular mini-chromosomes. We developed these approaches initially using the genome of *C. passerinae* 2 and then searched for similar patterns in the other three *Columbicola* taxa. Our methodology is further outlined in Figure 1.

**Figure 1.**
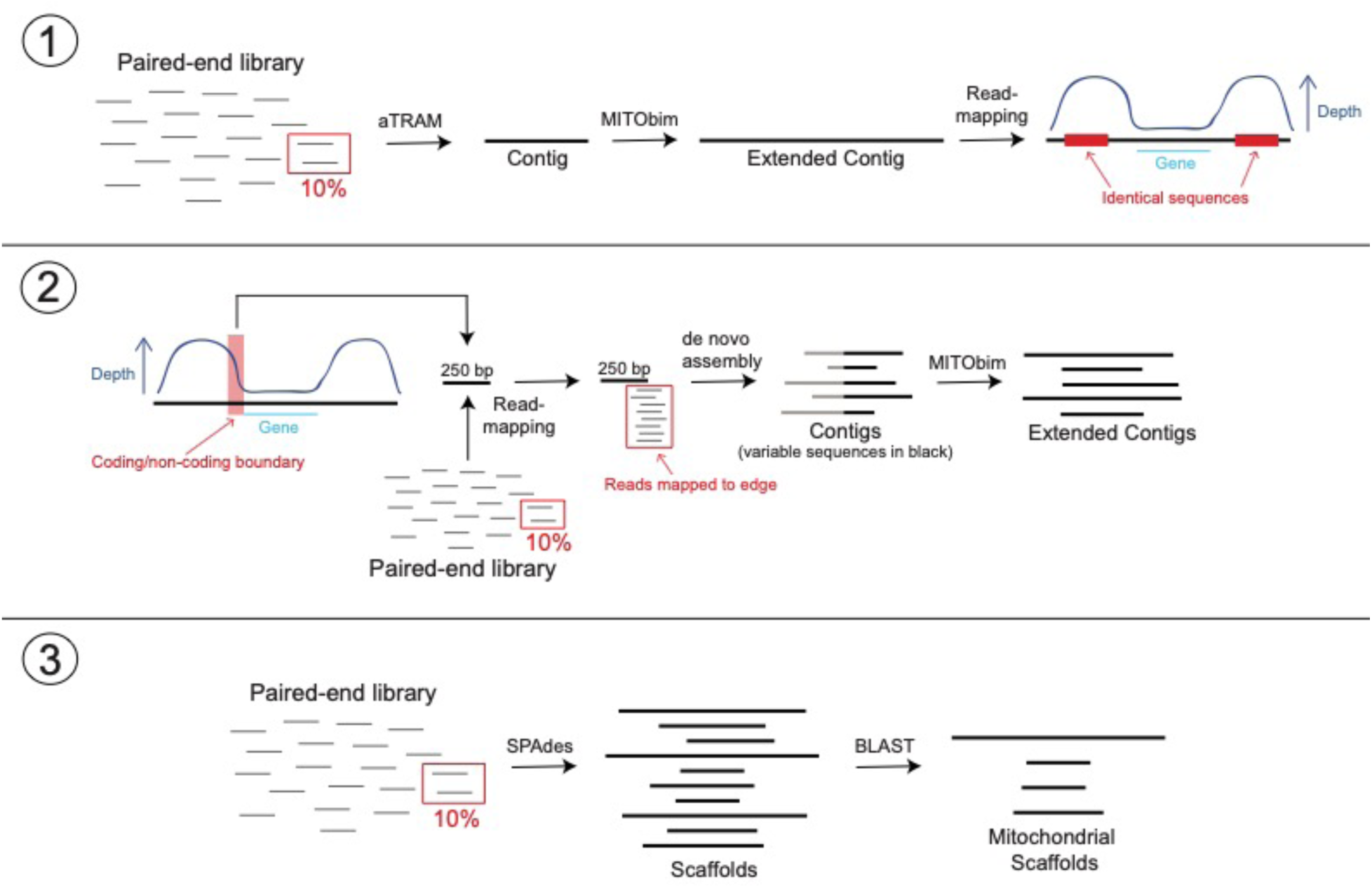
Workflow detailing three approaches for assembling the mitochondrial genomes of *Columbicola* lice. The first approach (1) uses de novo and read-mapping methods to assemble mitochondrial contigs and identify peaks in read coverage. The second approach (2) utilizes coverage information and MITOS annotations to identify the boundaries between coding and non-coding regions. Reads are then mapped to this boundary region. Reads that map to the 5’ edge are extracted, trimmed of conserved sequences, and used as starting references in MITObim. The third approach (3) is a more standard de novo assembly using SPAdes and identifying assembled mitochondrial scaffolds with BLAST.

First, we used a combination of aTRAM v.2.0 (Allen et al. 2018) and MITObim v.1.8 (Hahn et al. 2013) to assemble contigs of mitochondrial sequences. aTRAM uses BLAST (Altschul et al. 1990) to identify reads from a user-specified target and then uses third-party software to assemble those particular reads de novo. We attempted to assemble all 13 mitochondrial protein coding genes in *C. passerinae* 2 with aTRAM, using the amino acid sequences of each gene from the published *Campanulotes compar* mitochondrial genome as target sequences (AY968672; Covacin et al. 2006). For each assembly, we ran aTRAM for three iterations, used ABySS (Simpson et al. 2009) for de novo assembly, and used 10% of the read library. We used a fraction of the library because mitochondrial reads are often much more prevalent than nuclear reads in a shotgun sequencing library, and using only a portion of these reads helps to avoid erroneous assemblies associated with numts, highly-repeated motifs, or contaminants. We then used the best-scoring contig from each gene assembly as a starting reference in MITObim. MITObim is a pipeline that uses MIRA v.4.0.2 (Chevreux et al., 1999) to iteratively map reads against a starting reference (e.g., a single gene) and then against subsequent contigs to potentially reconstruct whole mitochondrial genomes. For each starting reference, we ran MITObim for a maximum of 100 iterations with the --quick option and utilizing paired-end information. We once again used 10% of the reads. Our MITObim runs resulted in a single contig for each starting reference. We trimmed the 5’ and 3’ ends of contigs that had many IUPAC ambiguities, because this can indicate issues with the read mapping. We then used Geneious v.11.1.5 (Biomatters Ltd., Auckland, NZ) to de novo assemble the trimmed contigs, using “Medium Sensitivity” and the default parameters. Finally, we tested for evidence of circularization in every contig, either from MITObim or the Geneious de novo assembly, by identifying identical stretches of sequence >50 bp near the 3’ and 5’ ends using megablast in the BLAST web interface.

Our second approach for assembling the mitochondrial genome of *C. passerinae* 2 leveraged the conserved control region (CR) among potential minicircles. Because the CRs have been shown to have sequence blocks that are highly conserved among the minicircles found within each species of mammal-associated louse, we expected there to be many more reads that map to the non-coding CR in a given contig compared to those which map to genic regions. However, identifying large read-coverage differences across a contig can be used for more than just evidence for presence/absence of minicircles. We can use this coverage information for a single contig to identify the boundary between the non-coding control and genic regions, and then map reads to this “boundary region” to identify reads from multiple minicircles. Song et al. (2019) used a similar approach to assemble fragmented mitochondrial chromosomes in mammal lice. To do this, we identified potential CRs by aligning the MITObim contigs with MAFFT (Katoh et al. 2002) in Geneious (default parameters) and identifying conserved regions among the contigs (Figure S1). We identified coding regions using the beta version of MITOS2 to annotate the MITObim contigs (Bernt et al. 2013; http://mitos2.bioinf.uni-leipzig.de/index.py). Using each annotated contig as a reference, we then mapped the paired-end reads used by MITObim for the assembly of that particular contig. Read-mapping was carried out in Geneious using “Medium-Low Sensitivity” and up to 5 iterations. As predicted, we identified coverage spikes upstream and downstream from the regions with annotated genes (Figure S2, Table 1). For the *cox1* contig, we extracted a 250 bp region immediately upstream from the coding region, and mapped the entire 10% paired-end read library (i.e., not only reads that mapped to the contig in MITObim) to this region using Geneious (same parameters as above). We then identified reads that mapped to and extended beyond the 3’ end of the reference. Theoretically, these reads include sequences from all minicircles. We assembled the reads de novo in Geneious and identified contigs with mean coverage >20X. We then aligned these contigs with MAFFT, trimmed the conserved 5’ ends of each contig, and used the trimmed contigs (which are therefore putative coding portions of the minicircles) as starting references in MITObim (max 100 iterations, 10% read library). We used BLAST to test for circularization in these resulting contigs and to identify identical regions of sequence from the conserved non-coding CRs.

**Table 1.** Read-mapping statistics for *Columbicola* and *Pediculus humanus* mitochondrial contigs assembled with MITObim using aTRAM-assembled starting references. Values are averaged across all contigs for each species. Statistics for each contig are listed in Table S1.

|  | Number of reads | Contig length | High coverage | Low coverage | Difference |
| --- | --- | --- | --- | --- | --- |
| *C. columbae* | 50501 | 7100 | 3034 | 747 | 2287 |
| *C. macrourae* | 17433 | 3177 | 1082 | 133 | 948 |
| *C. passerinae* 1 | 12155 | 2847 | 821 | 112 | 709 |
| *C. passerinae* 2 | 61459 | 3219 | 7315 | 665 | 6650 |
| *P. humanus* | 16251 | 3014 | 425 | 36 | 390 |

Third, we used a standard de novo approach to assemble the *C. passerinae* 2 mitochondrial genome. We used SPAdes v.3.11.1 (Bankevich et al. 2012) to assemble the 10% read library using the --plasmid option to invoke plasmidSPAdes (Antipov et al. 2016). We assessed the assembly quality with the web version of QUAST (Gurevich et al. 2013). We then converted the resulting scaffolds to a BLAST-formatted database in Geneious, and identified scaffolds containing mitochondrial protein-coding genes using *tblastn* with amino acid sequences from *Campanulotes compar* as queries.

Assemblies with aTRAM, MITObim, and plasmidSPADES were run on a cluster with 24 cores, two 2.60GHz Sky Lake processors, and 384 GB of memory maintained by Information Technology at Purdue (West Lafayette, IN, USA). De novo assembly, read mapping, and BLAST analyses in Geneious were run locally on a 2.9 GHz Intel i5 processor with 8 GB of memory.

To annotate the *C. passerinae* 2 mitochondrial genome, we initially used MITOS2 to annotate the contigs constructed from reads that mapped to the control/coding region boundary (approach #2). We then manually fine-tuned gene annotations (following the criteria in Cameron 2014b) and identified the number of minicircles by focusing on annotated regions with conserved flanking regions. We identified conserved regions by using BLAST to find highly similar sequences (>80% identical, e-value <1e^-20^) and searching for these sequences both within and between different contigs. We used the “boundary contigs” for annotation because the aTRAM and de novo approaches did not assemble all protein-coding genes. Nevertheless, to confirm all three approaches produced similar results, we aligned corresponding contigs with MAFFT in Geneious. We considered contigs “corresponding” if they shared protein-coding genes identified through MITOS/manual annotation (aTRAM and boundary contigs) or BLAST searches (de novo scaffolds). All three iterative assembly methods produced regions of identical sequence at the 5’ and 3’ ends of the contigs corresponding to the conserved areas of the CR shared by each minicircle in *C. passerinae* 2. Assemblies beyond this point were unreliable, as they can represent pooling of reads derived from multiple minicircles. MITObim assigns ambiguity codes to sites with alternative bases (similar to calling heterozygous alleles in nuclear data), which in this case likely indicates the mapping of reads from multiple minicircles. To account for this uncertainty, we retained the ambiguous calls of MITObim as “Ns” within the CRs. The identical region downstream of the genic region was trimmed off as this represents circularization.

After assembling and annotating the mitochondrial genome for *C. passerinae* 2, we wanted to search for similar patterns in our other three representatives of the *Columbicola* genus: *C. columbae*, *C. macrourae,* and *C. passerinae* 1. Evidence for minicircles in these taxa would be evidence that the architecture is conserved across the genus, and further support the validity of our approach to assembling the mitochondrial genome of *C. passerinae* 2. We used a similar approach with these other three taxa: first assembling longer contigs with aTRAM and MITObim, and then de novo assembling reads associated with the CR/coding region boundary to produce starting references for MITObim. We tested for a consistent architecture within *Columbicola* by using BLAST to compare our annotated *C. passerinae* 2 contigs to contigs from the other three species. We then used BLAST to search for conserved stretches of sequence flanking coding regions identified by MITOS and trimmed the CR assemblies as described above. We compared the variation in gene-order and content within mincircles between species and verified MITOS annotations following the process outlined in Cameron (2014b).

### Mitochondrial genome assembly of *Pediculus humanus*

As a confirmation of our bioinformatic approaches, we assembled the mitochondrial genome of *Pediculus humanus*, a species with known minicircles verified with both PCR and Southern Blotting (Shao et al. 2009). We downloaded the *P. humanus* A 2013 paired-end Illumina data from the GenBank SRA database (SRR5088471), cleaned the library, and assembled the minicircles using the same three approaches used for *Columbicola* (MITObim from aTRAM-assembled starting seeds, MITObim from CR/PCG edge starting seeds, and *de novo* with SPAdes). We used published gene sequences (translated PCGs and rRNA) from *P. humanus* (Shao et al. 2009) as targets for aTRAM. We then mapped reads from MITObim to the *cox1* assembly to identify a CR/protein-coding boundary and assemble contigs from reads that overlapped the boundary. Because the *Pediculus* library was considerable larger than the *Columbicola* libraries (128,102,260 cleaned reads), we used 1% of the library (compared to 10% of the *Columbicola* libraries) in each assembly approach.

## RESULTS

### Genome data

Paired-end library sizes varied among the four *Columbicola* genomes. There was an average of 7,598,374 reads from 10% of the adapter and quality-trimmed libraries. The number of reads ranged from 3,557,924 in *C. passerinae* 1 to 7,617,990 in *C. macrourae* (Table 2). We report statistics for 10% of the libraries because these were the reads used for assembling mitochondrial genomes.

**Table 2.** Mitochondrial genome assembly information for four *Columbicola* species.

|  | **Paired-end reads\*** | **SRA Accession** | **GenBank Accessions** | **Edge contigs** | **Minicircles** | **AT%** |
| --- | --- | --- | --- | --- | --- | --- |
| *C. columbae* | 7608708 | Pending | Pending | 19 | 15<sup>†</sup> | 68.3 |
| *C. macrourae* | 7617990 | SRR3161953 | Pending | 19 | 16 | 59.7 |
| *C. passerinae 1* | 3557924 | SRR3161931 | Pending | 17 | 17 | 65.1 |
| *C. passerinae 2* | 4553898 | SRR3161930 | Pending | 19 | 17 | 64.3 |

\* 10% of total reads
<sup>†</sup> Two protein-coding genes not assembled

### Mitochondrial genome assembly of *Columbicola passerinae* 2

The software aTRAM produced assembled contigs for seven protein-coding genes in *C. passerinae* 2 using the *Campanulotes compar* amino acid bait sequences: *cox1* (1,530 bp), *cox2* (873 bp), *cox3* (801 bp), *cob* (1,089 bp), *nad1* (783 bp), *nad3* (477 bp), and *nad5* (945 bp). The reported contig lengths are from the best-scoring contigs that we subsequently used as starting references in MITObim. Resulting MITObim contigs had the following lengths: *cox1*: 3,246 bp (5 iterations); *cox2*: 2,665 bp (7 iterations); *cox3*: 3,597 bp (10 iterations); *cob*: 3,673 bp (11 iterations); *nad1*: 2,973 bp (7 iterations); *nad3*: 2,465 bp (7 iterations); *nad5*: 3,913 bp (10 iterations). An average of 61,459 reads mapped to each contig in the final MITObim iteration (Table S1). When we mapped these reads to the MITObim contigs in Geneious, all contigs showed elevated coverage near the 5’ and 3’ ends of the contigs and lower coverage toward the centers of the contigs. However, coverages toward the centers were still high. For example, coverage at the ends (positions 1-805 and 2,378-3,246) of the contig produced with the *cox1* reference was 12 times higher than coverage toward the center (positions 806-2,377) of the contig (Figure S1; Table S1). The other contigs showed a similar pattern; among all contigs the coverage at the ends of each contig were on average 11 times higher than coverage near the centers of the contigs (Table 1, Table S1). When we annotated the contigs with MITOS, in every instance the lower-coverage regions corresponded to coding genes, whereas the regions with high-coverage peaks did not have any genes annotated with high confidence. We were unable to produce any additional contigs by assembling all the MITObim contigs together in Geneious. However, aligning each contig to itself with BLAST indicated >50 bp regions of identical sequences in the high-coverage 5’ and 3’ regions, thus suggesting each contig is a complete circle (Figure S1).

Although our first approach indicated there are several complete minicircles in *Columbicola*, we were not able to assemble every mitochondrial gene with this approach. One goal of our second approach was to assemble a set of contigs that contained all of the expected mitochondrial genes. The alignment of MITObim contigs from our first approach showed similar sequences near the ends of all contigs but highly variable sequences towards the middle of the contigs (Figure S2). The variable regions also corresponded to annotated genes and lower coverage. Together, these results suggest the conserved, high-coverage regions are part of the conserved CRs in the minicircles. We identified 6,400 reads from the 10% read pool (6,226,688 reads) that mapped to the boundary of a CR and the 5’ end of *cox1*. De novo assembly of these reads resulted in 19 contigs with ≥20X coverage. Alignments of the 19 contigs showed conserved sequences near the 5’ ends and variable sequences towards the 3’ ends (Figure S3). After trimming the conserved regions, the 19 contigs had an average length of 118 bp (min 31, max 139). Using the trimmed 19 contigs as starting references, MITObim produced contigs with an average length of 2,983 bp (min 1,814; max 4,213) from an average of 10 iterations per contig. BLAST identified >50 bp regions of identical sequences on the 5’ and 3’ ends of each MITObim contig.

The plasmidSPADES de novo assembly of the 10% read pool produced 2,024 contigs (N50: 12,832) and 2,020 scaffolds. We found BLAST hits for every *Campanulotes compar* protein-coding and ribosomal RNA gene among the scaffolds. However, *atp8*, *nad4l*, and *nad6* BLAST returns had high e-values >0.01 and low bit scores <25, indicating the homology of these sequences should be interpreted with caution. The longest scaffold with a high-scoring BLAST hit was 1,733 bp. These scaffolds were highly similar to corresponding contigs (i.e., containing the same protein-coding gene) assembled with MITObim (from both aTRAM and control edge starting references), with the only differences coming near or outside the gene boundaries identified by MITOS (Figure S4). However, none of the de novo-assembled scaffolds covered the entire length of a MITObim contig. Relevant data for all three assembly approaches, including starting references and resulting contigs/scaffolds, are available in Dryad (to be submitted upon acceptance).

### Annotation of the *Columbicola passerinae* 2 mitochondrial genome

An initial automated annotation of the control edge MITObim contigs with MITOS and subsequent manual annotations identified all 13 protein-coding genes, both rRNA genes, and 19 of the 22 tRNAs. These genes are separated on 17 separate mini-chromosomes. Although we recovered 19 contigs from MITObim, the assembly process can produce multiple contigs which possess identical coding regions, but whose CRs have sequence differences in portions of the CR outside of those conserved across all minicircles of *C. passerinae* 2. These ‘duplicate’ contigs were treated as assembly artefacts and removed from subsequent analyses. We determined the number of distinct chromosomes using a combination of the results described above, identifying conserved stretches of sequence and annotating the intervening regions (Figure S5). Each protein-coding and rRNA gene is on its own circular chromosome, with two chromosomes containing only tRNA genes. The chromosomes range from 1271 bp to 2862 bp with two contigs (*cox1* and *nad4*) forming incomplete circles, as their assemblies lacked non-coding sequence in the 3’ direction of the coding gene (Figure 2).

**Figure 2.**
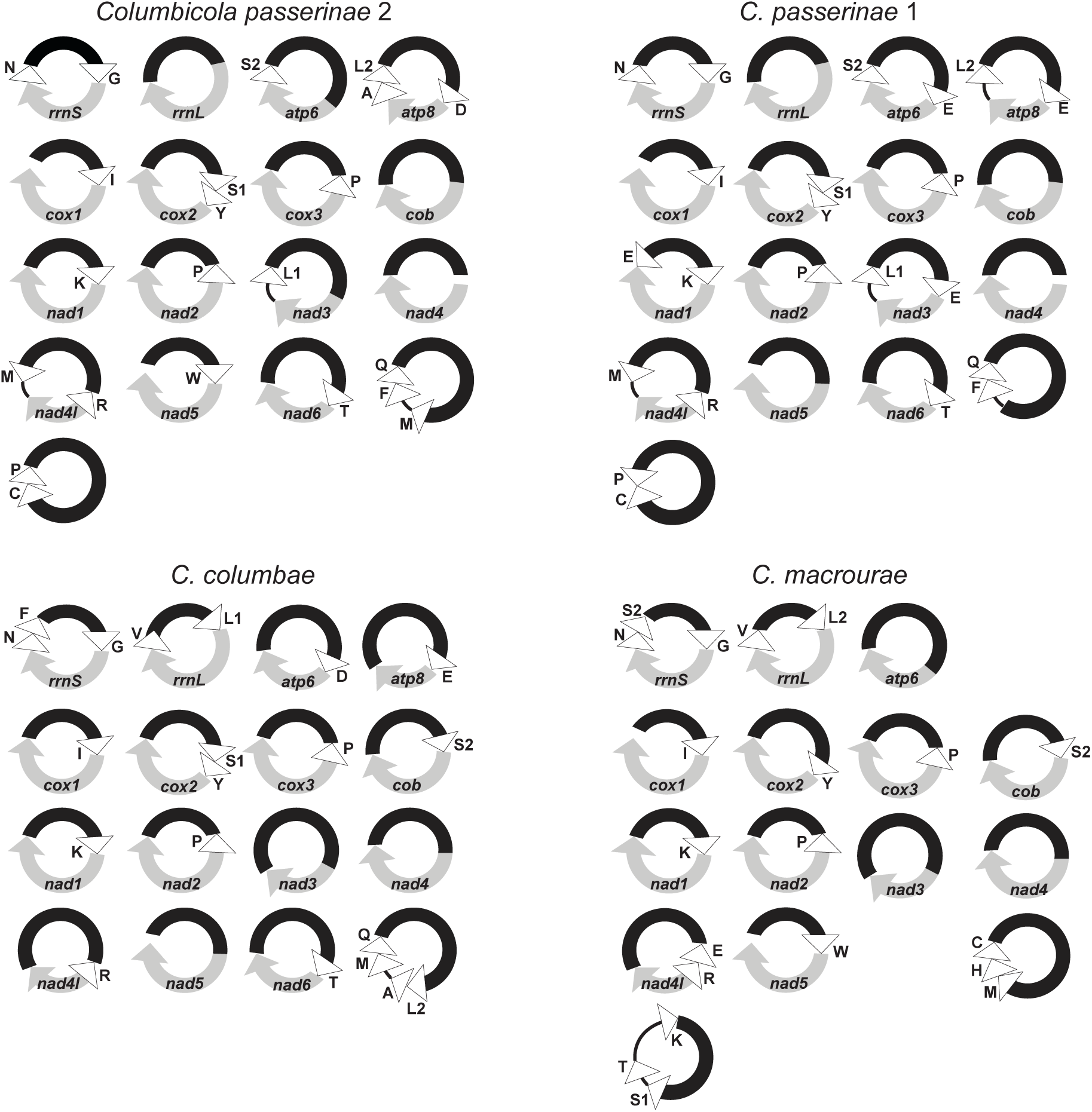
Mitochondrial genomes of four species of *Columbicola* lice: *C. passerinae* 2, *C. passerinae* 1, *C. columbae*, and *C. macourae*. Protein-coding and rRNA genes are indicated in gray, with arrows indicating 5’ to 3’ directionality in protein-coding genes. Annotated tRNAs are indicated with white triangles on the diagrams and are labeled with IUPAC amino acid letter abbreviations. Control regions are indicated with black segments. Partially assembled chromosomes have white space at the unassembled ends. Black lines indicate intergenic regions.

### Assembly of other *Columbicola* species

Assemblies of the three other *Columbicola* species showed similar patterns to *C. passerinae* 2. Contigs from MITObim using the aTRAM-assembled references did not assemble together in Geneious, but when aligned showed conserved regions of sequence with highly variable intervening regions. Although there was variation in the number of mapped reads and coverage values, for each species the conserved non-coding CRs in each contig had higher (4 to 9 times) coverage than the variable coding regions (Table 1). MITOS only identified genes in the variable, lower coverage regions. De novo assembly of reads that mapped to control/coding boundaries resulted in a similar number of contigs for most species (Table 2). MITOS was able to recover most genes from the resulting extended contigs from MITObim. BLAST searches identified similar regions flanking the annotated genes. As with *C. passerinae* 2, we used these flanking regions to identify individual minicircles. We recovered evidence for 15-17 minicircles in each species, with lengths of recovered complete chromosomes ranging from 1207-3118 bp for *C. passerinae* 1, 1262-3173 bp for *C. macourae,* and 1119-2657 bp for *C. columbae* (Figure 2).

### Assembly of *Pediculus humanus*

Assemblies of the *P. humanus* mitochondrial chromosomes showed similar patterns to the *Columbicola* assemblies. aTRAM assembled 11/13 PCGs (all excluding *nad3* and *nad4l*). Using these assemblies as starting references in MITObim produced contigs with an average length of 3,014 bp. Read-mapping to these MITObim contigs showed large relative coverage difference between coding and non-coding regions (average of 12 times higher coverage in non-coding regions; Table1 and Table SS). Using the contigs assembled from the CR/*cox1* edge reads, we recovered 17/18 chromosomes (we were unable to assemble the chromosome that contains *rrnS*) that showed high sequence similarity (>99% in coding regions) to the published *P. humanus* mitochondrial sequences from Shao et al. (2009) (Figure S6; contigs and alignments to published sequences are available in Dryad (to be submitted upon acceptance)). There was lower similarity between the non-coding CRs, but most of these contigs still had >90% total similarity to the published sequences. Four contigs had particularly lower similarity (70-80%) with published sequences in the CRs, and five contigs were shorter than their comparable published chromosomes. We were able to assemble and identify 11 PCGs and both rRNA genes from *de novo* assembly. However, as with *Columbicola*, these contigs were shorter than contigs produced from the other two assembly approaches; most assemblies seemed to fail at the CRs (Figure S7). There was also a chimeric contig containing portions of *nad5* and *rrnL*.

### Comparative mitochondrial genome structure among *Columbicola*

Overall mitochondrial genome structure was consistent across *Columbicola* species, with each protein-coding and rRNA gene on a separate chromosome and two minicircles with only tRNAs. All four genomes showed AT compositional bias (ranging from 59.7-68.3 AT%; Table 2), which is consistent with other insect mitochondrial genomes (Cameron 2014a). Additionally, protein coding genes aligned to the corresponding genes from the *C. passerinae* 2 assembly and had expected uncorrected distances between overlapping regions (13% between *C. passerinae* 1, and 22%-28% between the more distantly related taxa). None of the completely assembled genes were significantly longer or shorter than homologous copies in other *Columbicola* species or other louse species, which suggests the assemblies were reliable. However, *nad5* was incompletely assembled in *C. passerinae* 1 and *C. columbicola*. Assembled contigs from *C. macrourae* did not include minicircles with either *atp8* or *nad6*; however, this is likely an assembly or annotation failure rather than evolutionary loss of these genes within this species. Both genes are comparatively short and have higher rates of amino-acid substitution than other louse mt genes creating challenges for BLAST. Additional aTRAM and BLAST searches using the *C. passerinae* 2 amino acid sequences of *atp8* and *nad6* as references likewise failed to identify these two genes in *C. macrourae*, further highlighting the difficulty of assembling or annotating them from short-read genomic data. Similarly, each *Columbicola* species assembly was missing between two (*C. columbiae*) and five (*C. macrourae*) tRNA isotypes and had one or more duplicated tRNA isotypes. None of the canonical 22 tRNA isotypes were missing across all four *Columbicola* species, suggesting that missing tRNAs are likely present on other tRNA-only minicircles which failed to assemble or be adequately annotated.

Although the overall genome structure was consistent across the four *Columbicola* mitochondrial genomes, there was substantial variation in gene arrangement between *Columbicola* species. Of the 17 minicircles identified, only one (*trnI*-*cox1*) had a common gene arrangement across all four *Columbicola* species. Each of the other 14 minicircles containing a PCG or rRNA gene differed between at least two species due to the presence, absence, or translocation within the minicircle of accompanying tRNAs. These differences varied among comparisons: *C. passerinae* 2 and *C. passerinae* 1 shared 10 minicircles, *C. passerinae* 2 and *C. columbae* shared 7, *C. passerinae* 2 and *C. macrourae* shared 6, *C. passerinae* 1 and *C. columbae* shared 7, *C. passerinae* 1 and *C. macrourae* shared 4, and *C. macrourae* and *C. columbae* shared 9. The assembled and annotated chromosomes for all four *Columbicola* species are available on GenBank (submission upon acceptance). Relevant data generated from the various analyses, including assembled contigs, read pools, and read-mapping references, are available on Dryad (to be submitted upon acceptance).

## DISCUSSION

Fragmented mitochondrial genomes are a rare phenomenon in nature, with only a few animals known to possess this genomic architecture. Extensive fragmentation of the mitochondrial genome into a large number of minicircles each with a small number of genes per chromosome is even rarer still. Amongst the insects for which complete/near-complete mitochondrial genomes have been sequenced, highly fragmented, minicircle-type genomes have only previously been found in a clade of parasitic mammal lice (Anoplura, Rhynchopthirina, and Trichodectidae), while less fragmented genomes consisting of 2-3 chromosomes have been found in free-living booklice of the genus *Liposcelis* (Shi et al. 2016), and a species of thrips (Dickey et al. 2015). In insects for which partial mitochondrial genomes have been sequenced, evidence for genome fragmentation has also been recovered for several genera of avian feather lice (*Anaticola, Philopterus,* and *Quadraceps*) (Cameron et al. 2011). These exemplar chromosomes did not include the full suite of genes, although it is very likely the missing genes are present on other chromosomes that were not sequenced in these taxa. In the current study, we found strong evidence for highly-fragmented, minicircle-type genomes in another feather louse (Philopteridae) genus, *Columbicola*. We were able to assemble and annotate contigs which contained all of the protein-coding and rRNA genes in three species and 13 of the 15 genes in a fourth species (*C. macrourae*). We found no evidence for pseudogenes or for different copies of PCGs or rRNA genes on multiple minicircles, as has been found in the fragmented mitochondrial genome of the nematode *Globodera* (Gibson et al. 2007). We were also able to recover 17 to 20 of the 22 tRNA genes in each species, and all 22 isotypes in at least one species. Other reported minicircular genomes from sucking lice are also missing between one to two tRNA genes per species, consistent with our findings here (Song et al. 2019). It is possible the missing genes are on one or more additional minicircles that we could not detect with our approach. Nevertheless, with at least 17 minicircles and an average of two genes per circle (and as few as one), the *Columbicola* mitochondrial genome appears to be much more fragmented than that of any other feather louse species reported by Cameron et al. (2011), which had between 6 and 22 full genes on the recovered chromosomes. The mitochondrial genomes of *Columbicola* are also more fragmented than the mitochondrial genomes of most mammal associated lice (Jiang et al. 2013, Dong et al. 2014a, b, Herd et al. 2015, Shao et al. 2015, 2017). *Columbicola* has a similar level of fragmentation to the human louse (*Pediculus humanus*), which has at least 20 minicircles and an average of two genes per circle (Shao et al. 2009, 2012). Like *Columbicola*, *Pediculus* also has several circles that only contain tRNA genes. Although this genome organization has not been found in most other louse taxa with minicircles, in which all tRNAs that were detected are co-located with PCGs or rRNAs. However, undetected tRNA-only minicircles are a plausible explanation for putatively missing tRNA isotypes in these species.

Every recovered protein-coding gene in each species of *Columbicola* was on a separate minicircle. This is the first case of such an extreme level of fragmentation among PCGs in a metazoan mitochondrial genome. Even *Pediculus humanus*, which has a comparably fragmented genome in terms of the number of minicircles, has a circle containing both *atp6* and *atp8*. The genes *atp6* and *atp8* are consistently adjacent to one another in other fragmented or single louse mitochondrial genomes and have traditionally been understood as being co-transcribed genes (Senior 1988, Pelissier et al. 1992). The two genes are separated in the mitochondrial genome of the ibis louse *Ibidoecus*, but this species’ mitochondrial genome is not fragmented and has a canonical single mitochondrial chromosome (Cameron et al. 2011). In the case of *Columbicola* species, these genes appear to have separated onto different chromosomes. Once any fragmentation occurs, it is possible that there is no constraint on which parts of the genome become fragmented, because transcription already needs to be coordinated among different chromosomes.

There are considerable differences in tRNA gene arrangements among *Columbicola* species. Only one chromosome (*trnI*-*cox1*) was consistent across the four *Columbicola* species, and there are varying levels of shared arrangements between pairs of species. Degrees of similarity in minicircle gene arrangements appears to mirror phylogenetic relatedness: *C. passerinae* 2 and *C. passerinae* 1 share the most minicircles (13), whereas *C. passerinae* 2 shares just 7 with *C. columbae* and 6 with *C. macrourae*. *Columbicola passerinae* 1 and 2 are very closely related, being cryptic species, whereas *C. columbae* and *C. macrouae* are in two different clades (Boyd et al. 2017). This consistency among taxa suggests mitochondrial gene arrangements could help clarify phylogenetic relationships within *Columbicola* or aid in species delimitation within species complexes (e.g., *C. passerinae*) (Cameron et al. 2006, Cameron 2014).

Arrangement comparisons are, however, complicated by tRNA duplications which are present in all four *Columbicola* species. For example, *C. passerinae* 1 has four, sequence identical copies of *trnE* which result in three minicircles differing in gene content from *C. passerinae* 2, suggesting recent duplication of this gene into what are otherwise conserved minicircles. Other tRNA duplications appear themselves to be part of conserved minicircle gene arrangements. For example, *trnP* is found in minicircles with *cox3* and *nad2* in all four species, and in a tRNA-only minicircle with *trnC* in both *C. passerinae 2* and *C. passerinae* 1. Within each species, all copies of *trnP* are sequence identical with the exception of *C. columbae* where the *nad2* associated copy has a large insertion between the anticodon and acceptor stems disrupting the variable loop. The presence of sequence-identical, duplicate copies of tRNAs across multiple minicircles, combined with the variation in tRNA arrangement between *Columbicola* species is consistent with recombination (or other gene conversion mechanisms) between minicircles as has been proposed elsewhere (Shao et al. 2012; Dong et al. 2014b).

Overall, variation in which minicircles are present among different species of *Columbicola* is considerably higher than has been recorded for most other lice with minicircular mitochondrial genomes. Variation within a genus ranges from just 1 of 18 minicircles being different between two species of *Pediculus* (Herd et al. 2015) to 9 of 11 minicircles being different between species of *Polyplax* (Dong et al. 2014b). Other than *Polyplax*, over two-thirds of minicircles are shared by all members of a genus of sucking louse (17/19 to 4/12) suggesting that minicircle arrangements are stable at the genus-level within mammal-associated lice (Dong et al. 2014a). As might be expected given this level of variation, few derived gene-boundaries are shared between *Columbicola* species and other lice, of which only *trnI-cox1* and *trnY-cox2* are likely of phylogenetic significance being shared by the common ancestor of a clade comprising all Ischnocera, Anoplura, and Rhychophthirina (c.f. Shao et al. 2017).

### Assembling fragmented mitochondrial genomes with a fully bioinformatic approach

Assembling fragmented mitochondrial genomes can be a challenging and labor-intensive process. In this study, we were able to assemble the mitochondrial minicircles of lice using a fully bioinformatic approach from raw genomic Illumina reads. In particular, we used an approach that assembled short, variable contigs from reads that mapped to the edge of the control region (CR) and coding regions. These contigs served as starting references for assembling longer contigs, essentially acting as “bioinformatic PCR primers.” By identifying conserved non-coding CRs that flank adjacent coding regions, we could then detect separate chromosomes and confirm circularization of each contig. Subsequent annotations recovered most of the 37 mitochondrial genes present in metazoans.

We used two other methods for assembling mitochondrial genomes: *de novo* and read-mapping extension. *De novo* assembly is fast (∼12 minutes on five cores for SPAdes) and has the advantage of not requiring a starting reference. However, finding mitochondrial scaffolds is reliant on BLAST searches with a closely related target. In our case, we were not able to reliably identify every PCG or rRNA gene in each *Columbicola* species, presumably because the initial target was too distantly related (*Campanulotes*), the missing genes were particularly divergent (e.g., *nad6*), or the genes simply failed to assemble. It is also challenging to identify tRNA-only minicircles with a *de novo* approach, because tRNA genes are short (<100 bp) and can be highly divergent from even a closely-related target. Our *de novo* assemblies also had trouble assembling through the CR because of the high similarity of this region among minicircles, and as a result the *de novo* scaffolds are considerably shorter than contigs produced from other approaches (Figure S4). Assemblies with aTRAM were also fast (assembled PCGs in ∼3.5 minutes total) and using the assembled genes as starting seeds with read-mapping in MITObim allowed us to extend the contigs into the CR. However, we were limited by the starting seeds we could assemble with aTRAM, which as with standard *de novo* assembly tended to exclude faster-evolving PCGs and tRNA genes. Our approach based on the coding/non-coding boundary combines the benefits of *de novo* and read-mapping approaches, while also enabling us to assemble all mitochondrial minicircles (perhaps excluding small chromosomes that only contain tRNAs).

Our approach is an important progression that builds on previous work that primarily used wet laboratory methods or a combination of bioinformatics and laboratory methods to identify and confirm mitochondrial minicircles. The first minicircular mitochondrial genome from insects, that of *Pediculus humanus*, was assembled in 18 separate contigs from Sanger shotgun reads, with non-coding regions sequenced from clone libraries of PCR products generated by designing primers within the coding regions of assembled contigs (Shao et al. 2009). Circularization was also confirmed by restriction digest and Southern hybridization of purified mitochondrial DNA extractions from *Pediculus* (Shao et al. 2009). We were able to recover these same chromosomes from *Pediculus* using our bioinformatic approaches. Subsequently, a procedure was developed for sequencing minicircles by first sequencing several partial coding gene sequences, designing outward facing primers which would amplify the non-coding portion of several minicircles, then designing primers in conserved regions of the non-coding region to amplify all minicircles present in a given species. This mixed population of amplicons would then be sequenced using NGS methods to yield clean, minicircle contigs (Shao et al. 2012). Recently, Song et al. (2019) simplified this approach to minimize primer design and PCR steps by using initial partial coding sequences (generated by universal PCR primers) as seeds to directly assemble minicircles from genomic extracts. Similar to our approach, conserved CRs were identified for select mitochondrial genes (initially sequenced with Sanger) from whole genome assemblies and subsequently used to identify all contigs derived from mitochondrial minicircles. Finally, Song et al. (2019) confirmed putative minicircle findings with PCR amplification and gel electrophoresis.

Approaches that rely on laboratory methods to either identify or confirm fragmented mitochondrial genomes are certainly useful and should continue as an option for these types of studies, particularly because there are some limitations to assembling these genomes with Next Generation Sequencing (NGS) data. For example, because it is repeated across multiple minicircles, it is challenging to assemble through long stretches of relatively conserved non-coding regions (e.g., CRs) with short-read data. The entire CR may not assemble for a particular minicircle, which prohibits confirmation of circularization. This was the case for a few of our assembled fragments (incomplete circles in Figure 2). Due to their similarity, reads from the CRs could also misassemble among different minicircles, so the exact sequences of the CR should be interpreted with caution or reported with ambiguity codes (as we have done in this study). Additionally, a bioinformatic approach may fail to recover all genes present in the organism. This could particularly be an issue for tRNA-only minicircles or highly divergent PCGs such as *atp8* and *nad6*. However, laboratory-based methods can also fail to identify minicircles or genes, especially tRNAs. Additionally, the most rigorous wet-lab methods for confirming the presence of minicircles, such as Southern blotting, require large volumes of starting tissue, thus limiting their utility to species that are either physically large (unlike lice) or are maintained in culture, which is rarely feasible for the vast majority of louse species.

It should also be noted that bioinformatics methods do not appear to erroneously assemble fragmented mitochondrial genomes for taxa where they are not present. Many hundreds of insect mitochondrial genomes have been assembled from NGS datasets (e.g. Linard et al. 2018) using bioinformatics assembly procedures such as those presented here (e.g. *de novo* or MITObim baited assembly). Complete circularized mitochondrial genomes are routinely recovered, or when incomplete, assemblies typically only lack portions of the CR (e.g. Gomez-Rodriguez et al. 2015). In these assemblies there is no evidence for chimeras between incompletely assembled portions of genomes. Within insects, evidence for mitochondrial fragmentation derived from NGS sequencing and bioinformatics analysis is confined to taxa for which prior evidence for fragmentation had been collected (such as lice). Conversely, even within taxa such as lice where aberrant mitochondrial genomes appear to be common, NGS-only approaches have successfully assembled complete, canonical, single-chromosome mitochondrial genomes when such an architecture was present in a given taxon (e.g. for *Falcolipeurus*: Song et al. 2019; for *Campanulotes*: *unpub. data*).

Thus despite potential limitations, with the an ever-increasing reliance on NGS data it is important to demonstrate that fragmented genomes can be assembled from sequence read libraries. In this regard there is capacity for the study of “aberrant” mitochondrial genome architectures to follow the path of “conventional” mitochondrial genome studies over the past decade where the expansion of NGS techniques has led to a sharply reduced reliance on PCR-centric methods (see review Cameron 2014a). Avoiding PCR is especially relevant for rare specimens or for incorporating publicly available data (NCBI SRA), where it may not be possible to confirm genomic architecture with PCR due to a lack of DNA extracts. We assembled mitochondrial fragments from whole genome shotgun sequencing, but our approach could also be used with reads generated from target capture methods (e.g., ultra-conserved elements (UCEs)). These genome reduction methods target nuclear regions, but mitochondrial reads are also captured at relatively high depth as “by-catch” in the process (Amaral et al. 2015, Ströher et al. 2017). Utilizing the ever-growing diversity of genomic data, most of which is publicly available through databases such as GenBank, allows for further exploration of mitochondrial genome architecture and arrangements on a broad taxonomic scale. The bioinformatic methods described here accurately reconstruct aberrant mitochondrial genomes whose structures have been verified by wet-lab procedures (i.e. *Pediculus*) and extend our capacity to search for fragmented mitochondrial genomes into taxa which cannot be verified by wet-lab methods (i.e. *Columbicola* and other feather-lice).

### Evolutionary implications for highly fragmented mitochondrial genomes in *Columbicola*

Highly fragmented mitochondrial genomes in lice were first identified in human lice (*Pediculus humanus*) (Shao et al. 2009), and minicircle-type mitochondrial genomes have since been confirmed in several other genera of sucking lice (Anoplura; Jiang et al. 2013, Dong et al. 2014a,b, Herd et al. 2015, Shao et al. 2017), elephant and warthog lice (Rhyncophthirina; Shao et al. 2015), and chewing lice in the family Trichodectidae (Cameron et al. 2011, Song et al. 2019). Phylogenetic analyses have indicated that these three lineages of mammalian lice form a monophyletic group, which suggests highly fragmented mitochondrial genomes evolved once in the common ancestor of this group, as suggested by Song et al. (2019). Given the rarity of this architecture in other groups of organisms and the implications of fragmentation for disease, it is very reasonable to conclude there have been few transitions from full to fragmented mitochondrial genomes. However, our study implies a second such transition to minicircle-type mitochondrial genomes in parasitic lice. The mitochondrial genomes of *Columbicola* species are the first recorded instance of a fragmented mitochondrial genome from within the family Philopteridae for which the complete genome has been sequenced. Other published complete mitochondrial genomes of Philopteridae include species of *Bothriometopus* (Cameron et al. 2007), *Ibidoecus* (Cameron et al. 2011), *Falcolipeurus* (Song et al. 2019), *Campanulotes* (Covacin et al. 2006), and *Coloceras* (Cameron et al. 2011). All five of these species have a single circular mitochondrial chromosome, although *Coloceras* has a full-sized chromosome and at least one additional heteroplasmic fragment missing approximately one third of the full-sized genome. Recent phylogenomic analysis shows that *Columbicola* is nested within these other three taxa (Johnson et al. 2018), so the minicircularized genomes of *Columbicola* likely represent an independent transition from a single chromosome (Figure 3). It is possible that fragmentation occurred multiple times within the *Columbicola* genus, which could explain why we recovered fewer minicircles in *C. columbae.* However, given the similar levels of fragmentation and conserved gene order in some minicircles among the four *Columbicola* species, fragmentation is at most an infrequent phenomenon within the genus and most parsimoniously represents a single independent origin. A more densely-sampled analysis is needed to confirm the history of fragmentation within the genus.

**Figure 3.**
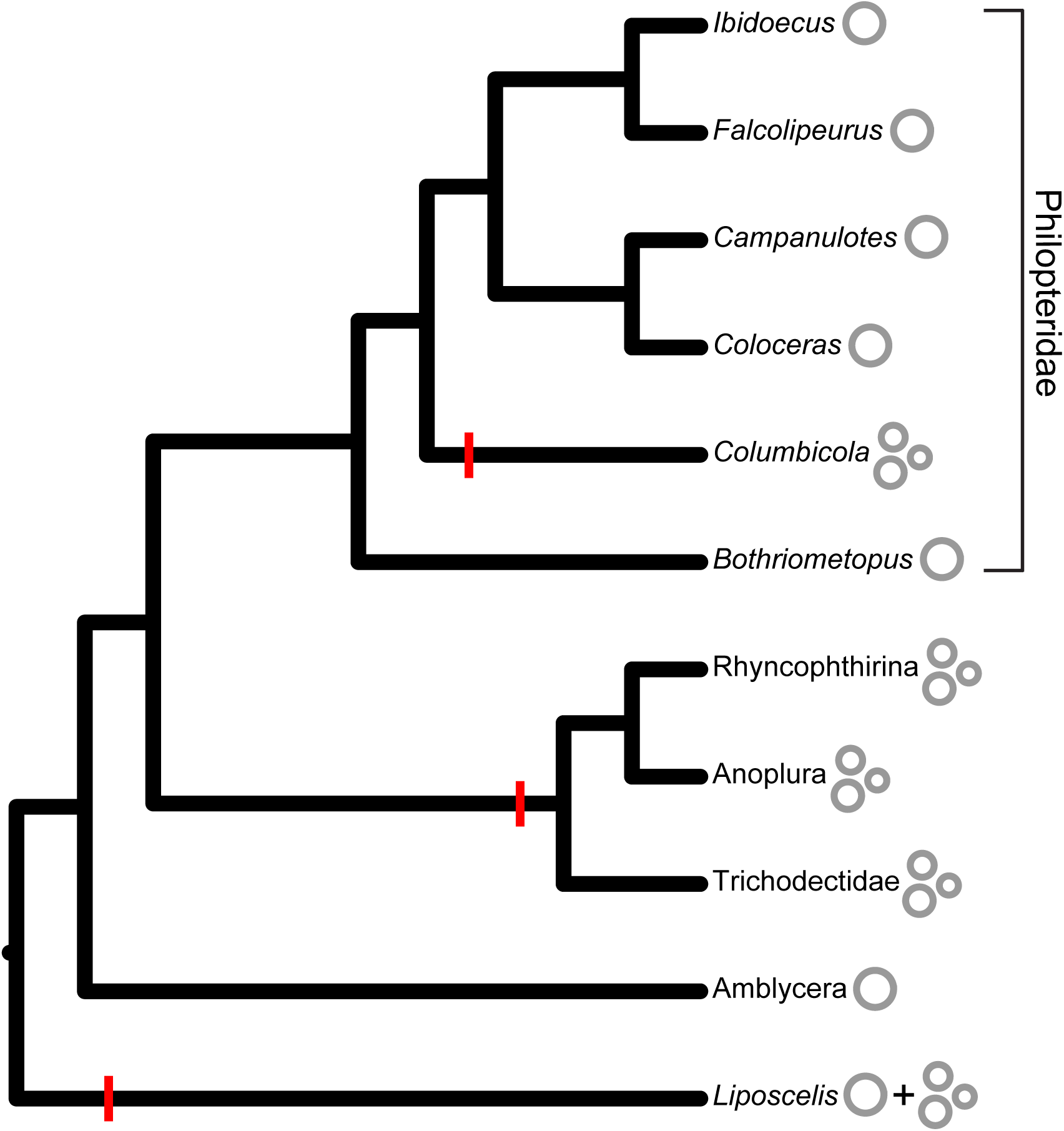
Cladogram that shows mitochondrial minicircles have evolved multiple times in parasitic lice and free-living relatives (*Liposcelis*). The phylogeny is adapted from Johnson et al. (2018) and features taxa with published full mitochondrial genomes, including *Columbicola* from the current study. Genome architecture (single or fragmented) is indicated next to the tip names. Presumed transitions from single to fragmented mitochondrial genomes are represented by red lines on the relevant branches.

Cameron et al. (2011) identified partial fragmented mitochondrial genomes in three other genera of Philopteridae (*Anaticola*, *Quadraceps*, and *Philopterus*), so it is possible other transitions to fragmented genomes from single chromosomes have occurred in these feather lice. However, none of these taxa possess the minicircle-type genome architecture (i.e., few genes per chromosome and a relatively long CR with conserved motifs) reported here for *Columbicola*. The independent origin of mitochondrial minicircles in *Columbicola* and mammal-associated lice allows for phylogenetically independent examination of the life-history traits proposed to be related to this phenomenon. As *Columbicola* are feather-feeding and associated with birds, the proposals that mitochondrial minicircularization in lice is related to blood- or mammal feeding are unlikely (c.f. Dong et al. 2014b). Similarly, the generation-time (24 days) of *Columbicola*, which is the most fragmented mitochondrial genome yet sequenced, falls within the range of generation times (Martin 1934) for other lice whose mitochondrial genomes are complete, or at least less fragmented, suggesting that genome fragmentation is not causally related to short generation times. Multiple independent transitions from single to fragmented mitochondrial chromosomes in one group of organisms (parasitic lice) do, however, suggest that there are underlying potentially shared mechanisms for the formation of minicircles, such as the lack of a functional mtSSB gene, and/or similar selective pressures retained across relatively deep evolutionary time (∼65 my). Future work can address these questions by building on the genome architectural framework established in this study.

## Conclusion

The results from this study demonstrate that highly-fragmented, mitochondrial minicircles are present in species of the feather louse genus *Columbicola*. Based on a well-supported phylogeny of parasitic lice, mitochondrial minicircles in *Columbicola* also imply that mitochondrial fragmentation originated multiple times in lice, which is surprising given the reported detrimental effects of fragmentation and how infrequently the phenomenon has been observed across Metazoa. These results should encourage further investigation of mitochondrial architecture in insects and other animals. Additionally, we were able to assemble the complete *Columbicola* mitochondrial genome from NGS data using a fully bioinformatic approach, thus contributing to a methodological trend that has had increasing momentum in the last decade. Applying this approach to *P. humanus*, a louse species with known mitochondrial minicircles, showed the method to be reliable and consistent with previous laboratory-based methods for confirming the presence of minicircles. As NGS data becomes more commonplace in sequenced-based studies, it is important to continue developing approaches that can extract useful information, particularly for organisms with challenging or uncommon features.

## ACKNOWLEDGMENTS

Alvaro Hernandez and Christ Wright at the Roy J. Carver Biotechnology Center (University of Illinois, Champaign, IL, USA) were instrumental in facilitating genome library preparation and sequencing. We also thank Jyothi Thimmapuram at the Purdue University Bioinformatics Core (West Lafayette, IN, USA) for helping to set up and run software on the Purdue cluster. This work was supported by the National Science Foundation [DEB-1239788 and DEB-1342604 to KPJ]; and the Australian Research Council [FT120100746 to SLC].

