## Supplemental material for "Mitochondrial genomes of *Columbicola* feather lice are highly fragmented, indicating repeated evolution of minicircle-type genomes in parasitic lice"

**Table S1.** Expanded read-mapping statistics for *Columbicola* and *Pediculus* mitochondrial contigs assembled with MITObim using aTRAM-assembled starting references. Each contig is listed according to the protein-coding gene used as a starting refence (e.g., *cox*1). Coverages are averages from particular regions (high or low coverage) of the contigs.

| | **Number of reads** | **Contig length** | **High coverage** | **Low coverage** | **Difference** |
|---|---|---|---|---|---|
| ***C. columbae*** | | | | | |
| *atp*6 | 38,041 | 4,379 | 4,191 | 696 | 3,495 |
| *cox*1 | 68,514 | 10,812 | 1,921 | 552 | 1,369 |
| *cox*2 | 52,254 | 7,133 | 3,732 | 717 | 3,015 |
| *cox*3 | 44,465 | 5,119 | 3,768 | 1,018 | 2,750 |
| *cob* | 51,501 | 7,586 | 2,749 | 930 | 1,819 |
| *nad*1 | 45,419 | 6,550 | 2,854 | 796 | 2,058 |
| *nad*4 | 61,401 | 9,311 | 2,296 | 750 | 1,546 |
| *nad*5 | 42,409 | 5,912 | 2,761 | 515 | 2,246 |
| ***C. macrourae*** | | | | | |
| *cox*1 | 17,362 | 3,572 | 925 | 148 | 777 |
| *cox*2 | 17,074 | 3,084 | 1,151 | 138 | 1,013 |
| *cox*3 | 17,278 | 3,365 | 1,034 | 170 | 864 |
| *cob* | 17,199 | 3,016 | 1,105 | 151 | 954 |
| *nad*1 | 18,902 | 3,043 | 1,155 | 149 | 1,006 |
| *nad3* | 16,208 | 2,396 | 1,310 | 96 | 1,214 |
| *nad5* | 18,007 | 3,763 | 891 | 82 | 809 |
| ***C. passerinae 1*** | | | | | |
| *cox*1 | 11,834 | 4,288 | 579 | 139 | 440 |
| *cox*2 | 13,504 | 2,538 | 873 | 119 | 754 |

| | | | | | |
|---|---|---|---|---|---|
| *cox*3 | 11,246 | 2,356 | 954 | 127 | 827 |
| *cob* | 13,449 | 2,913 | 795 | 108 | 687 |
| *nad*1 | 9,363 | 2,385 | 831 | 89 | 742 |
| *nad*3 | 12,923 | 2,172 | 916 | 128 | 788 |
| *nad*5 | 12,764 | 3,280 | 800 | 75 | 725 |
| ***C. passerinae 2*** | | | | | |
| *cox*1 | 53,383 | 3,246 | 7,582 | 636 | 6,946 |
| *cox*2 | 61,562 | 2,665 | 8,674 | 899 | 7,775 |
| *cox*3 | 70,512 | 3,597 | 7,272 | 515 | 6,757 |
| *cob* | 66,200 | 3,673 | 5,930 | 599 | 5,331 |
| *nad*1 | 55,295 | 2,973 | 7,951 | 642 | 7,309 |
| *nad*3 | 62,137 | 2,465 | 7,546 | 724 | 6,822 |
| *nad*5 | 61,121 | 3,913 | 6,249 | 639 | 5,610 |
| ***Pediculus humanus*** | | | | | |
| *atp6* | 16,348 | 2,687 | 485 | 39 | 446 |
| *atp8* | 16,338 | 3,238 | 426 | 39 | 387 |
| *cox1* | 16,656 | 3,306 | 406 | 40 | 366 |
| *cox2* | 16,490 | 2,827 | 492 | 34 | 458 |
| *cox3* | 15,840 | 2,756 | 408 | 31 | 377 |
| *cob* | 15,648 | 2,668 | 407 | 36 | 371 |
| *nad1* | 15,994 | 2,965 | 377 | 27 | 350 |
| *nad2* | 16,322 | 3,080 | 452 | 40 | 412 |
| *nad4* | 16,600 | 3,474 | 407 | 41 | 366 |
| *nad5* | 16,123 | 3,161 | 340 | 34 | 306 |
| *nad6* | 16,404 | 2,991 | 479 | 33 | 446 |

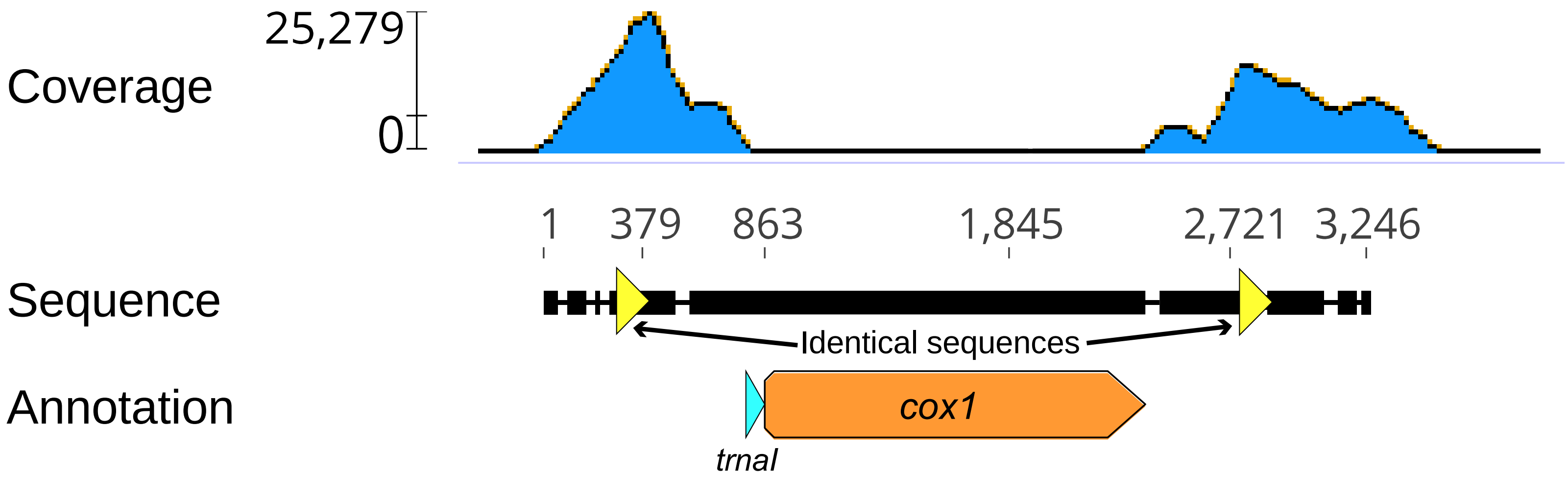

**Figure S1.** Graphical example of the coverage differences between coding and non-coding regions of *Columbicola* mitochondrial contigs. The contig was assembled in MITObim using *cox*1 from *C. passerinae* 2 (assembled from aTRAM) as a starting reference. Coding genes annotated from MITOS are indicated below the sequence. Regions of identical sequences are indicated with yellow triangles. Read coverage distribution was obtained by mapping reads used in MITObim to the assembled contig.

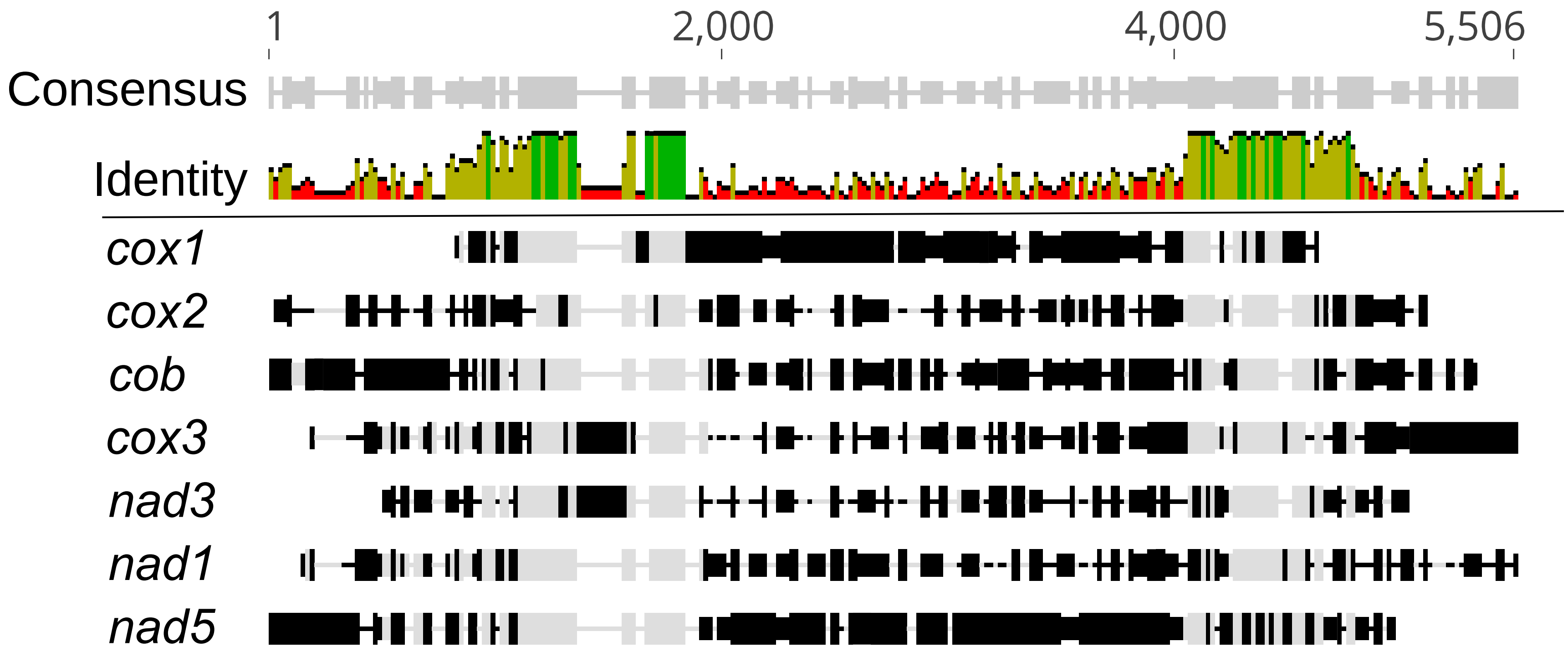

**Figure S2.** Multiple sequence alignment (from MAFFT) of MITObim contigs from *Columbicola passerinae* 2 produced using aTRAM-assembled starting references. Contigs are labeled according to the protein-coding gene used as a reference. Identical sequences among the contigs are indicated with gray and variable sequences with black.

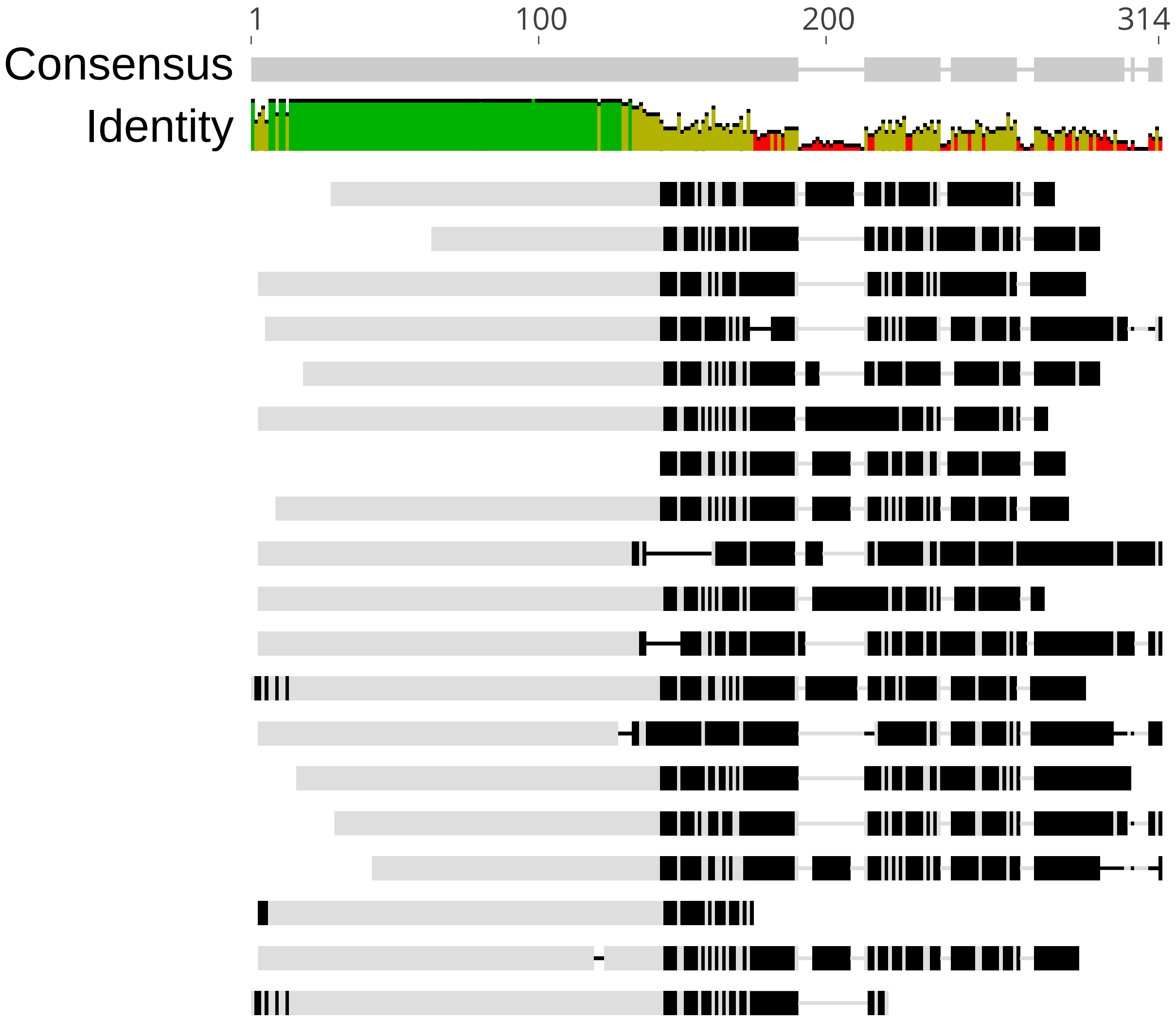

**Figure S3.** Multiple sequence alignment (from MAFFT) of 19 contigs assembled de novo from reads mapped to the edge of a 250 bp region between the boundary of a coding (*cox*1 and tRNA) and non-coding region in *Columbicola passerinae* 2. Identical sequences among contigs are indicated with gray and variable sequences with black. The variable regions were subsequently used as starting references in MITObim.

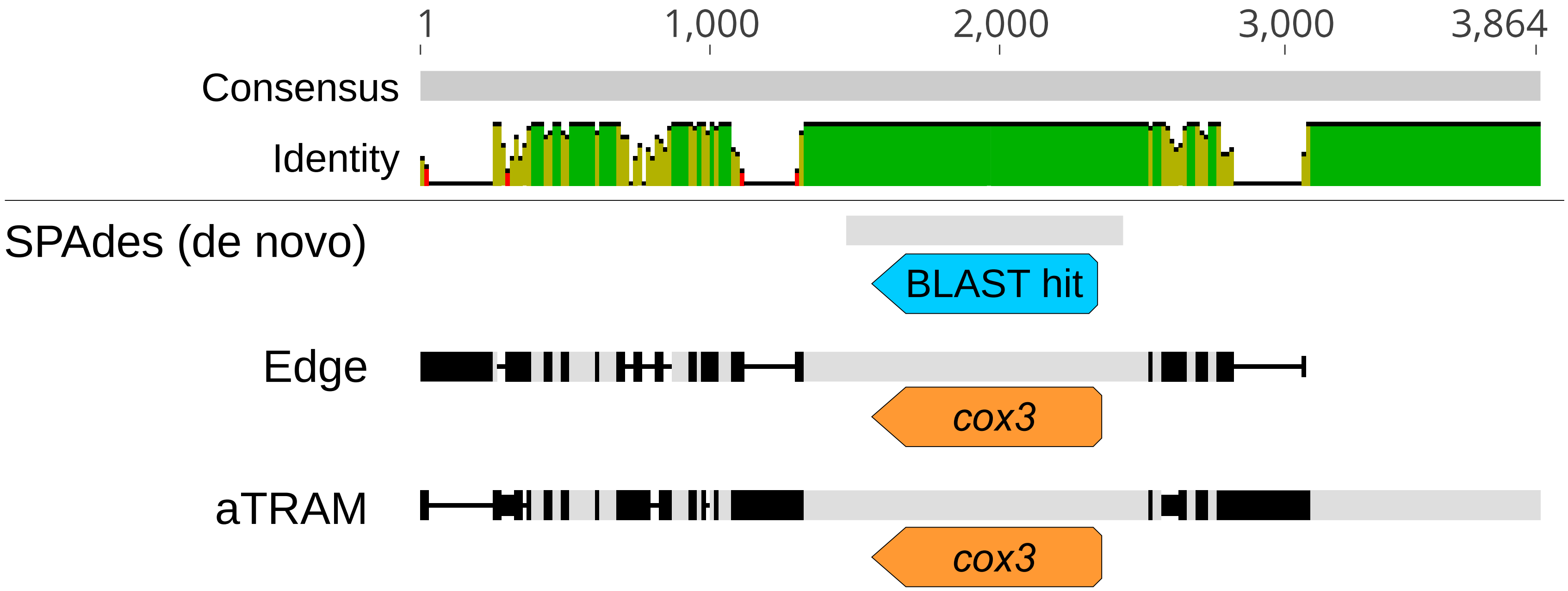

**Figure S4.** Example of an alignment (from MAFFT) of *Columbicola passerinae* 2 mitochondrial contigs produced through three different approaches: de novo, MITObim with "edge" starting references, and MITObim with aTRAM-assembled starting references. Gray indicates identical sequences among the contigs and black indicates variable sequences. Annotation from MITOS are shown below the Edge and aTRAM contigs (in this case *cox*3). The BLAST hit from searches with a *cox*3 query (*Campanulotes compar*) is shown below the de novo contig.

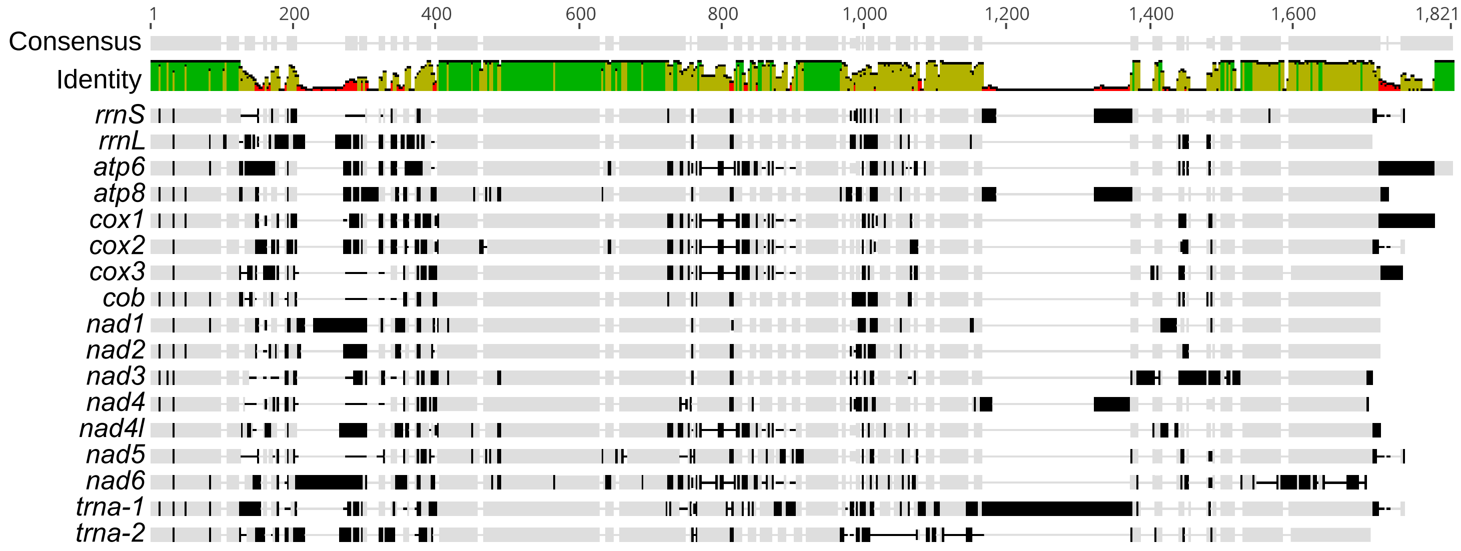

**Figure S5.** Alignment (from MAFFT) of non-coding control regions from each mitochondrial chromosome of *Columbina passerinae* 2. Names for each contig, labeled according to the gene content of each chromosome, are indicated to the left of the alignment. Gray indicates identical sequences among chromosomes and black indicates variable sequences.

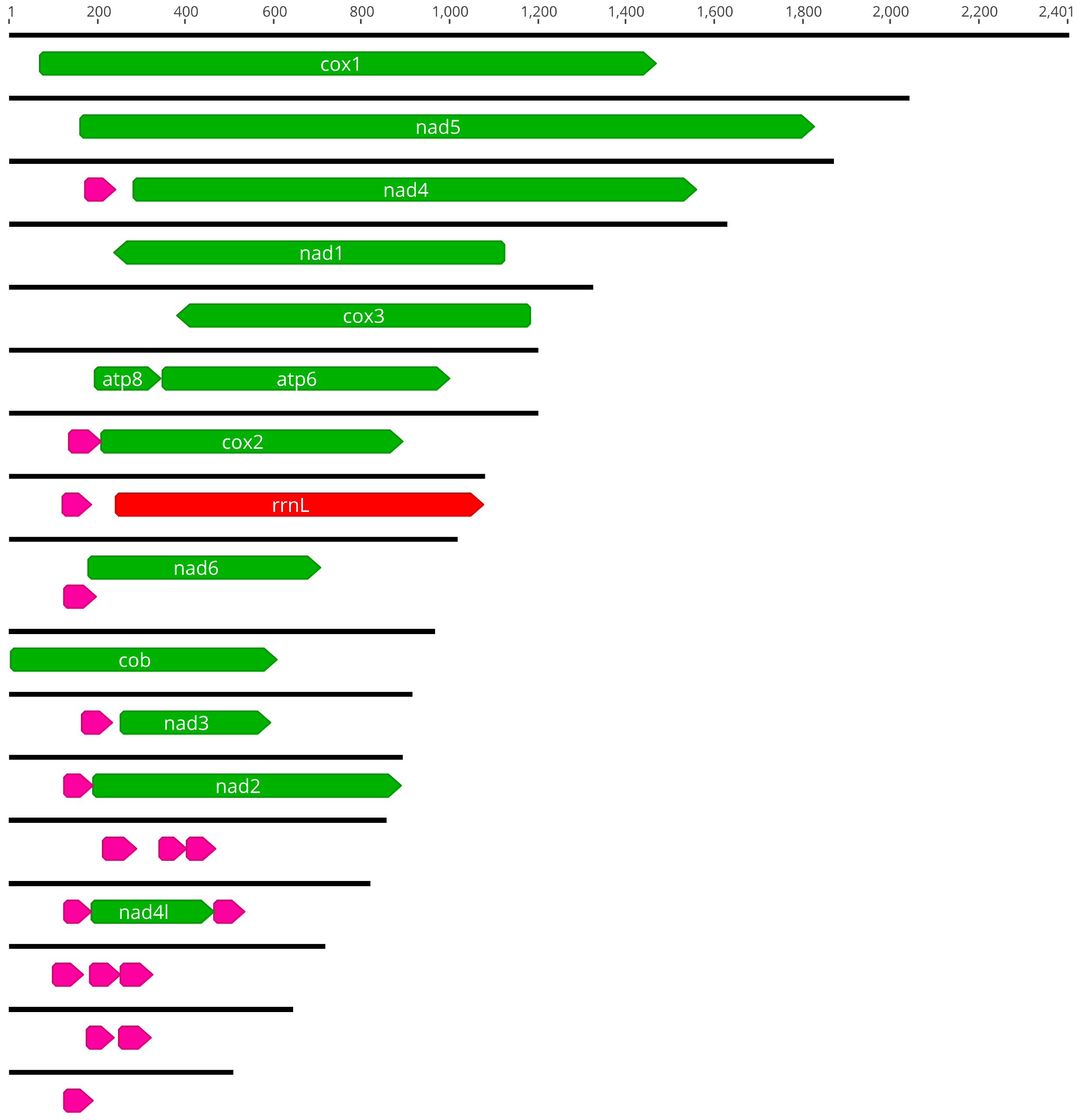

**Figure S6.** Mitochondrial contigs of *Pediculus humanus* assembled from reads that overlap the CR/*cox1* boundary. Each contig has been annotated with MITOS2 and trimmed according to published mitochondrial chromosomes of *P. humanus*. Protein-coding genes are indicated with green arrows, rRNA genes with red, and tRNA genes with pink. Lengths (bp) are indicated with the scale at the top of the figure.

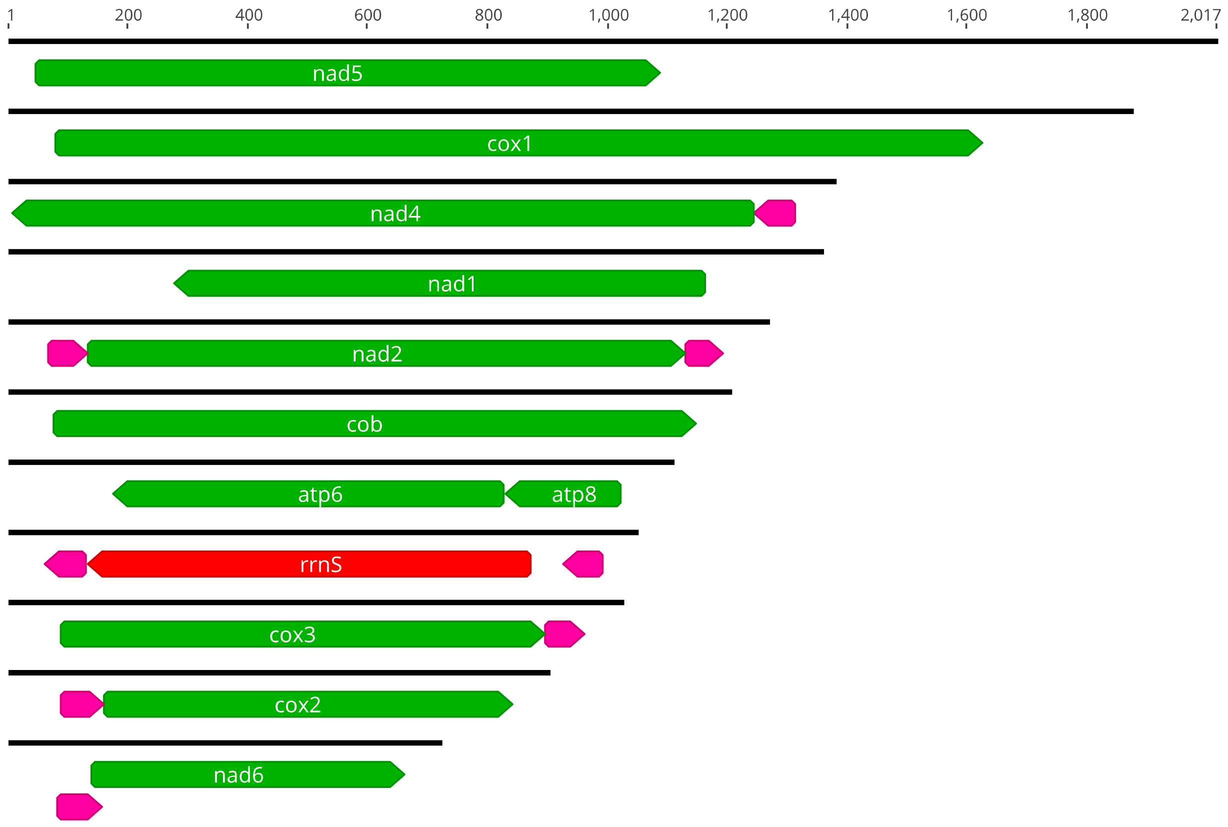

**Figure S7.** Mitochondrial contigs of *Pediculus humanus* assembled with a de novo approach in SPAdes. Each contig has been annotated with MITOS2. Protein-coding genes are indicated with green arrows, rRNA genes with red, and tRNA genes with pink. Lengths (bp) are indicated with the scale at the top of the figure.
